# Reversing an extracellular electron transfer pathway for electrode-driven NADH generation

**DOI:** 10.1101/476481

**Authors:** Nicholas M. Tefft, Michaela A. TerAvest

## Abstract

Microbial electrosynthesis is an emerging technology with the potential to simultaneously store renewably generated energy, fix carbon dioxide, and produce high-value organic compounds. However, limited understanding of the route of electrons into the cell remains an obstacle to developing a robust microbial electrosynthesis platform. To address this challenge, we engineered an inward electron transfer pathway in *Shewanella oneidensis* MR-1. The pathway uses native Mtr proteins to transfer electrons from an electrode to the inner membrane quinone pool. Subsequently, electrons are transferred from quinones to NAD^+^ by native NADH dehydrogenases. This reverse functioning of NADH dehydrogenases is thermodynamically unfavorable, therefore we have added a light-driven proton pump (proteorhodopsin) to generate proton-motive force to drive this activity. Finally, we use reduction of acetoin to 2,3-butanediol via a heterologous butanediol dehydrogenase (Bdh) as an electron sink. Bdh is an NADH-dependent enzyme, therefore, observation of acetoin reduction supports our hypothesis that cathodic electrons are transferred to intracellular NAD^+^. Multiple lines of evidence indicate proper functioning of the engineered electrosynthesis system: electron flux from the cathode is influenced by both light and acetoin availability; and 2,3-butanediol production is highest when both light and a poised electrode are present. Using a hydrogenase-deficient *S. oneidensis* background strain resulted in a stronger correlation between electron transfer and 2,3-butanediol production, suggesting that hydrogen production is an off-target electron sink in the wild-type background. This system represents a promising genetically engineered microbial electrosynthesis platform and will enable a new focus on synthesis of specific compounds using electrical energy.

## Introduction

Microbial electrosynthesis is a technology that utilizes microbes for production of useful chemicals using carbon dioxide, water, and electricity as feedstocks (1). While most microbial electrosynthesis efforts to date have targeted fuel production, other possible applications include production of platform chemicals or bioplastics (2). With pressure to develop sustainable production systems, microbial electrosynthesis is an attractive technology to simultaneously produce valuable products and store electricity generated by wind and solar technologies. Initial efforts have focused on three primary platforms for microbial electrosynthesis: pure cultures of acetogens (3, 4); undefined mixed cultures (5, 6); and electron shuttles paired with model bacterial strains (7, 8). While these approaches have demonstrated proof-of-concept, significant improvements in product yield and product spectrum are necessary to make microbial electrosynthesis economically viable.

Pure culture microbial electrosynthesis systems have been developed in an effort to understand the mechanism of electron uptake from a cathode. These systems primarily utilize acetogenic bacteria because they are naturally capable of converting carbon dioxide into acetate using hydrogen or other inorganic electron donors (9, 10). Multiple reports show that acetogens are capable of using an electrode in place of native electron donors (3, 4). While acetate is a relatively low value product, genetic modification may yield strains capable of producing other compounds through electrosynthesis (11). While this is a promising approach, significant challenges to acetogen engineering remain, and further developments are necessary (11, 12). An alternative approach that may yield specific products is to utilize chemical electron mediators to transfer electrons from an electrode into existing model organisms. This approach has been successful in multiple organisms to generate products such as succinate and ethanol (7, 13–15). However, the cost of the mediator and downstream separations likely make this approach too costly for industrial scale up.

Mixed culture approaches to microbial electrosynthesis have also borne success, due to their ability to generate a range of products without genetic modification. Early efforts in this area have primarily generated acetate, with adjustments to operating conditions driving increased yield (16–18). However, other molecules can also be produced, including butyrate, butanol, and ethanol (5, 19). Selective enrichment of the community is important to improve yields and product specificity (6, 17). While evidence that higher value products are attainable increases the attractiveness of electrosynthesis using undefined mixed cultures, off-target production remains problematic. For example, increased abundance of methanogens over time has posed difficulties for long term experiments due to decreased acetate titers and increased methane production (6, 20). Further, a mixed community cannot be engineered to produce a specific high-value compound with any current technology.

Overall, optimization of existing approaches to microbial electrosynthesis has been hampered by poorly defined interactions between bacteria and cathode electrodes. In most cases, the mechanism of electron transfer is unknown, although in mixed culture systems, H_2_ is recognized as a major electron carrier between the electrode and microbes (5). For pure cultures, direct electron transfer has been proposed, but not proven (3, 4). Similarly, specific mechanisms of electron transfer via chemical mediators are only beginning to be investigated (13). In contrast, transmembrane electron transfer is much better understood between dissimilatory metal reducing bacteria and anode electrodes. *Shewanella oneidensis* MR-1 is particularly well understood and the structure of the Mtr pathway and its function in transferring electrons to the outer surface of the cell have been thoroughly described in the last two decades (21–28).

The well understood electron transfer mechanism of *Shewanella* makes it an excellent candidate to engineer a microbial electrosynthesis system from the ground up using synthetic biology tools. Initial steps to explore the use of *S. oneidensis* MR-1 for microbial electrosynthesis have demonstrated reverse electron transfer through the Mtr pathway. Ross et al. (29) showed electron transfer from a cathode to *S. oneidensis* MR-1 with fumarate as an electron sink, indicating that the Mtr pathway can transfer electrons from a cathode to respiratory quinones. More recent work by Rowe et al. (30) has demonstrated that *S. oneidensis* MR-1 can also catalyze electron transfer from a cathode to oxygen through the Mtr pathway and the quinone pool.

While previous work has demonstrated the ability to generate reduced quinones via a cathode, this is not sufficient to power intracellular reduction reactions. Electrons in the quinone pool have too positive a redox potential to be spontaneously transferred to NAD^+^ (ca. −80 mV_SHE_ for menaquinone:menaquinol vs. ca. −320 mV for NAD^+^:NADH) and have only been transferred to electron acceptors with a more positive redox potential, such as fumarate (ca. +30 mV_SHE_). Electrons should ideally be transferred to an intracellular, lower potential electron carrier to be useful for electrosynthesis. NADH is a promising intracellular electron carrier to target because it is involved in many metabolic reactions and is relatively accessible from the quinone pool through NADH dehydrogenases. Under typical conditions, NADH dehydrogenases catalyze the favorable transfer of electrons from NADH to quinones and conserve energy as a proton-motive force (31). However, NADH dehydrogenases may also catalyze the reverse redox reaction, utilizing proton-motive force as an energy source, as observed in purple photosynthetic bacteria (32).

Based on previous work and our knowledge of *S. oneidensis* MR-1 metabolism, we hypothesized that NADH dehydrogenases could be driven in reverse to transfer electrode-derived electrons into the cell. Therefore, we engineered a system relying on the native Mtr pathway and NADH dehydrogenases to catalyze the electron transfer steps and using a heterologous light-driven proton pump to generate proton motive force (PMF) to drive NADH dehydrogenases in reverse (Figure 1). We expressed butanediol dehydrogenase and provided its substrate, acetoin, to provide an electron sink and a mechanism to track NADH generation. This system represents a generalizable platform to use cathodes to drive NADH-dependent reductions in *S. oneidensis* MR-1. Further development of this platform could lead to a new capability to generate specific, high-value compounds using electricity and carbon dioxide as feedstocks.

**Figure 1.**
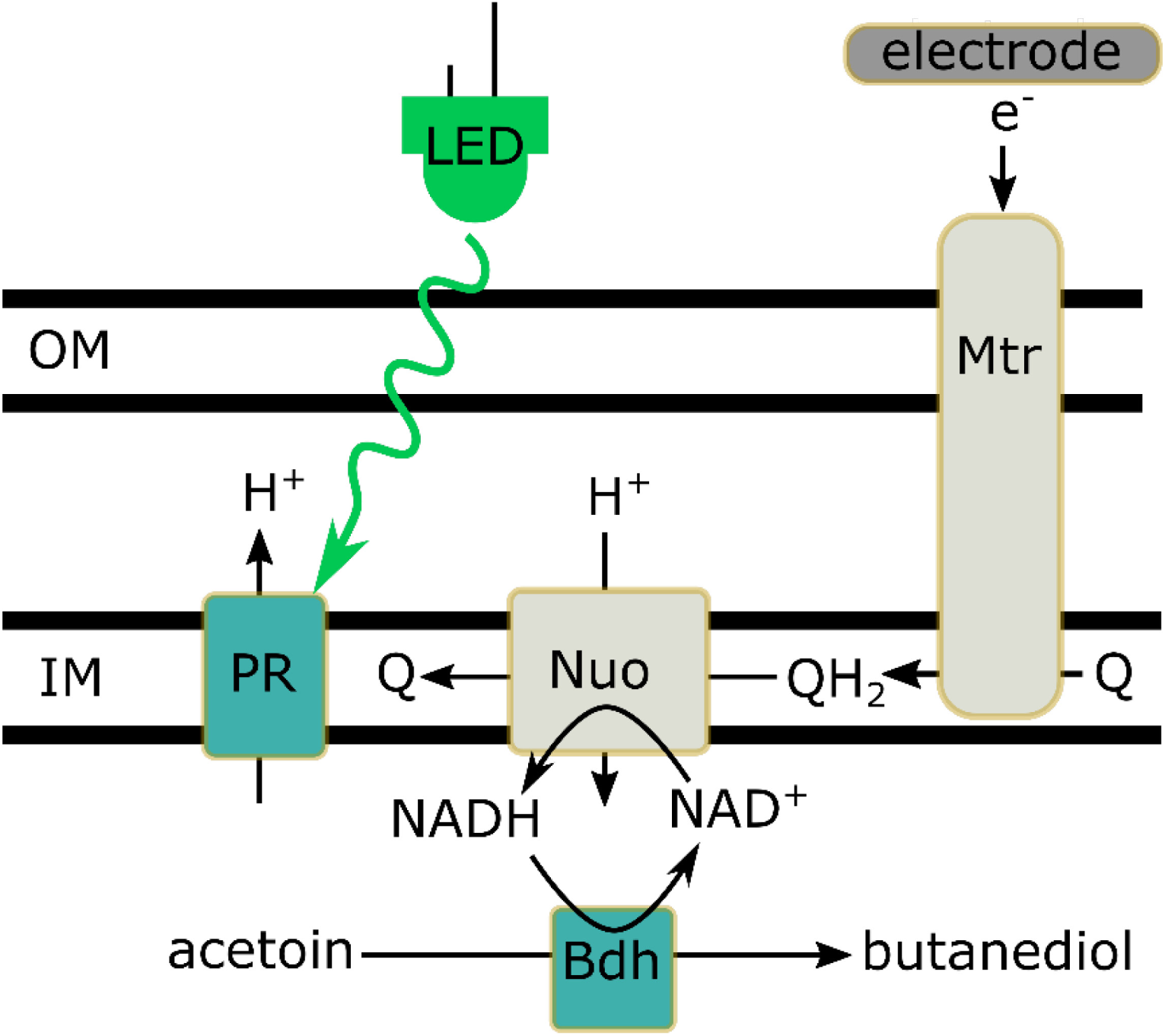
Engineered inward electron transfer pathway. *Shewanella oneidensis* MR-1 pathway designed to transfer electrons from the electrode to NAD^+^ to generate intracellular reducing equivalents. Native proteins are shown in grey, heterologous proteins are shown in green. OM: outer membrane; IM: inner membrane; LED: green light source. Nuo is one of four NADH dehydrogenases encoded in the *S. oneidensis* MR-1 genome.

## Results and Discussion

### Development of modified *S. oneidensis* strains

Genes encoding proteorhodopsin (PR) and butanediol dehydrogenase (Bdh) were cloned into a medium copy plasmid with a constitutive promoter and conjugated into *S. oneidensis* (both WT and a strain lacking hydrogenases; ∆*hyaB*∆*hydA*) (Table 1). A FLAG tag was added to the C-terminus of each protein to facilitate detection by Western blot. Both proteins were detectable in cleared lysates using an anti-FLAG antibody, confirming successful expression (**Figure S1**). To confirm function of Bdh, the modified strain was grown aerobically in the presence of 15 mM exogenous acetoin in LB. Accumulation of 2,3-butanediol was observed by HPLC, indicating that the expressed Bdh is functional (**Figure S2**). Function of PR expressed from a similar plasmid was previously demonstrated in *S. oneidensis* MR-1 (33, 34).

**Table 1.**
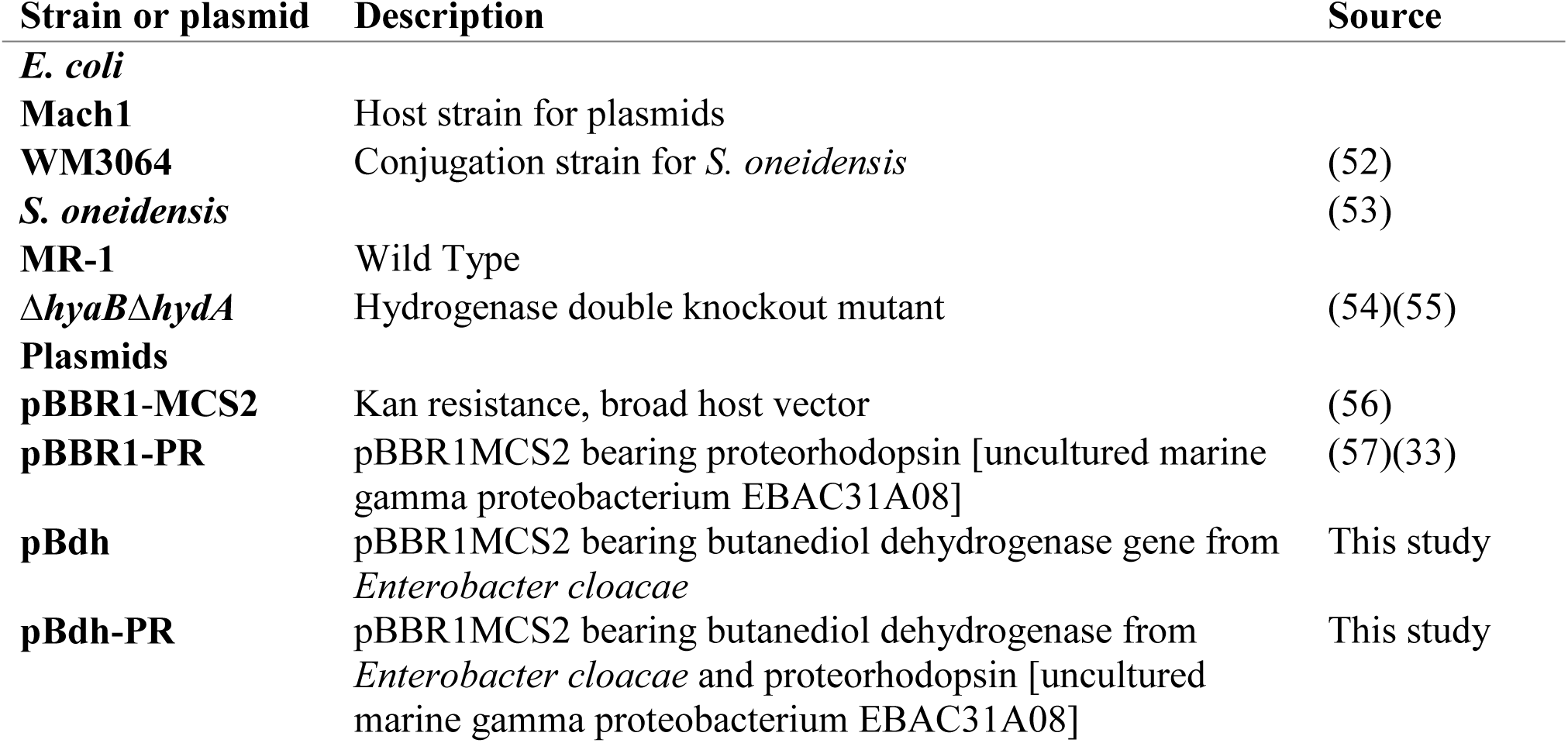
Strains and plasmids used in this study

### Light and current drive a target reduction reaction in modified *S. oneidensis*

To determine the capability of the engineered cells to accept electrons from a cathode, the strain lacking hydrogenases and carrying a plasmid with PR and Bdh was pre-grown and inoculated into bioelectrochemical systems at a final density of OD_600_ = 0.17. This mutant strain was used to ensure that H_2_ did not act as a mediator between cells and the electrode. This mutant was previously shown to lack the ability to produce or consume H_2_ (35, 36). To minimize the influence of organic carbon as an electron donor, the bioelectrochemical systems initially contained oxygen (ambient) and the working electrode was set at an anodic potential (+0.4 V_SHE_). This was done to decrease the availability of organic carbon by oxidizing stored organic material and media components carried over during inoculation. After 6 hours, the working electrode was switched to a cathodic potential (−0.3 V_SHE_), and N_2_ sparging was used to remove oxygen from the bioreactors. Each bioelectrochemical system was equipped with green LED lights around the working electrode chamber to drive proton pumping by PR (**Figure S3**). After switching to the working electrode to a cathodic potential, a stable cathodic current between −5 and −8 µA was observed. After 16 hours, an anoxic acetoin solution was injected into each bioreactor to a final concentration of 10 mM. After acetoin injection, an increase of −12 to −18 µA was observed in all bioelectrochemical systems (Figure 2). Note that for cathodic current, a greater negative current represents a greater amount of electron transfer.

**Figure 2.**
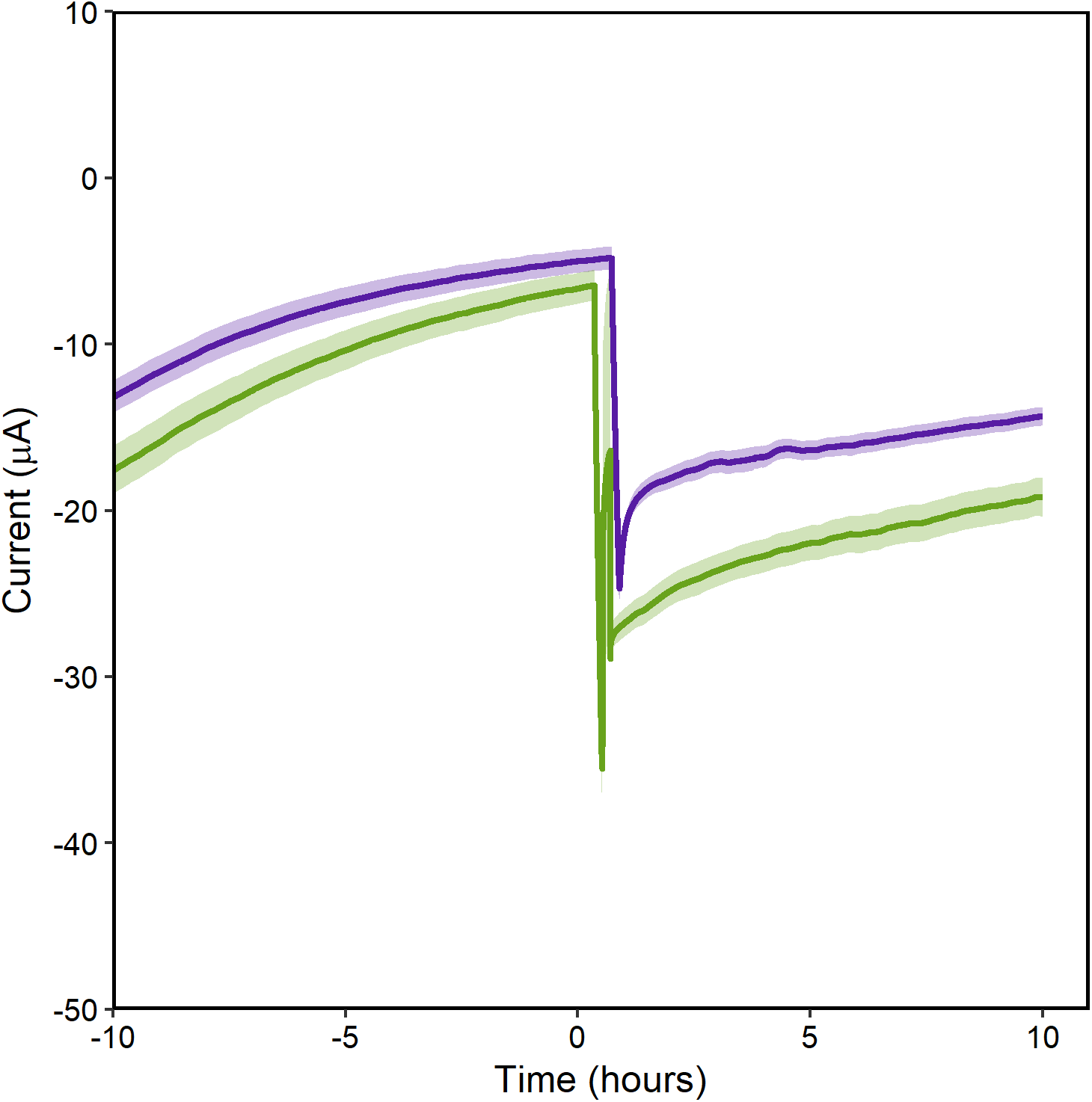
Cathodic current measurement during electron acceptor injection. Cathodic current increases significantly after the electron acceptor, acetoin, is added to the bioelectrochemical system. Time 0 represents acetoin injection. Current measured in bioelectrochemical systems containing cells grown with holo-PR are shown in green and those containing apo-PR are shown in purple. Each line represents the average of three replicates with standard error shown in transparent ribbons.

To determine the influence of proton pumping on electron uptake, we compared cells with and without functional PR by pre-growing the strain with or without the essential PR cofactor, all-*trans*-retinal. We refer to PR with retinal as holo-PR and PR without retinal as apo-PR. Cathodic current was significantly higher, (p=0.03) in systems with holo-PR (Figure 3). We confirmed that the increased current was due to PR activity by turning off the LED lights attached to the bioreactor. When the lights were turned off, cathodic current generated by cells with holo-PR decreased, while cells with apo-PR were unaffected (Figure 3). This supports the model that the effect of light is dependent on functional PR; if the effect of light was due to heating or interaction of light with native components, we would expect cells with apo-PR to have a similar response to cells with holo-PR. The dependence of current on holo-PR and light supports our hypothesis that enhanced PMF generation by PR promotes electron uptake by the modified strain.

**Figure 3.**
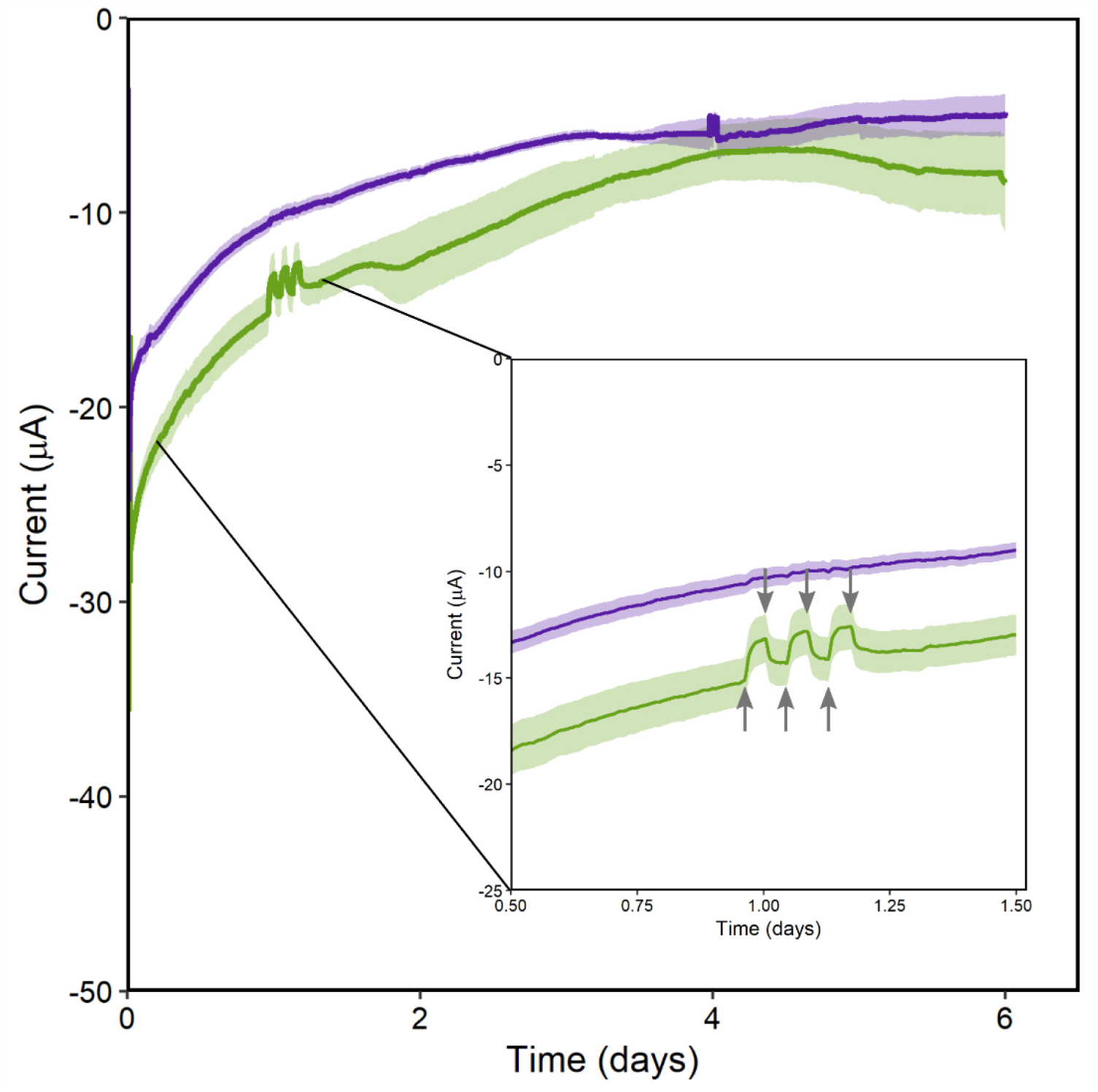
Response of cathodic current to green light. Current was measured in bioelectrochemical systems with electrodes poised at −0.3 V_SHE_. Current shown was measured beginning at injection of acetoin (time 0). Current measured in reactors containing cells with holo-PR are shown in green and those containing apo-PR are shown in purple. Inset graph shows current change due to removal or addition of green light. Lights were turned off for 3 one hour intervals beginning just before time = 1 day. Each line represents the average of three replicates with standard error shown in transparent ribbons.

During the same experiment, we also measured 2,3-butanediol accumulation to determine whether cathodic electrons were directed toward the target reaction, acetoin reduction. As with current, 2,3-butanediol accumulation was greater in the bioelectrochemical systems with holo-PR (Figure 4). Based on the total charge transfer over the course of 6 days, accumulation of 0.17 ± 0.02 mM 2,3-butanediol was expected by the end of the experiment. Accumulation of 0.19 ± 0.01 mM 2,3-butanediol was actually observed. The experiment was repeated using the same conditions, with the exception that the electrode was disconnected from the potentiostat. This experiment confirmed that acetoin reduction was driven by the electrode (Figure 4). Indeed, 2,3-butanediol production was significantly reduced when the electrode was not poised (p=0.01), and accumulation of only 0.13 ± 0.00 mM was observed. Based on our observations, 0.06 mM of the 2,3-butanediol (32% of total production) was generated through the electrode-dependent process. We hypothesize that 2,3-butanediol was not completely eliminated when the electrode was not poised because remaining organic carbon from the inoculum could be oxidized to generate the NADH needed for acetoin reduction. Although we attempted to remove as much organic carbon from the experiment as possible, some cells may have lysed during the washing and inoculation process, thus providing organics to the surviving cells.

**Figure 4.**
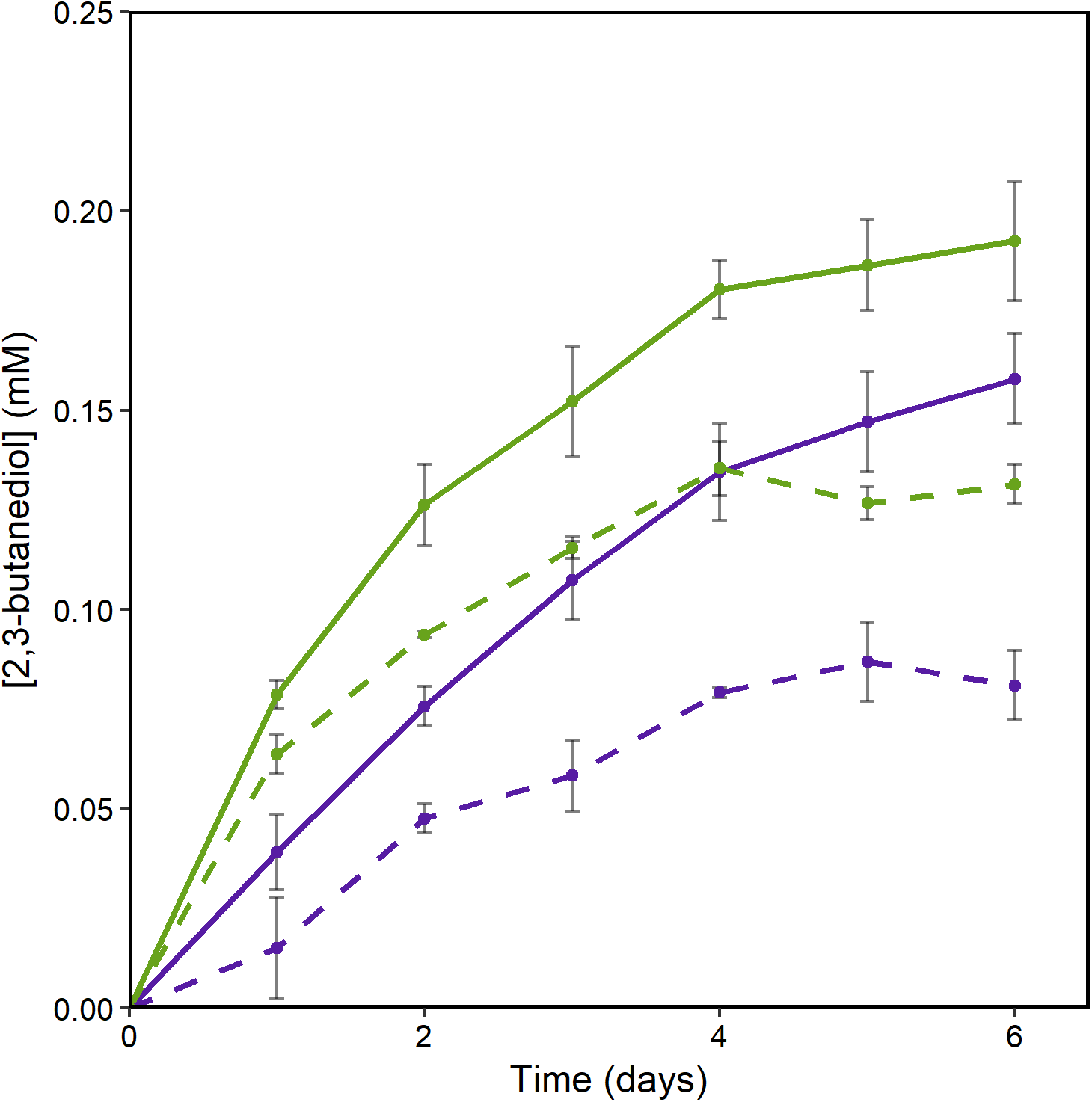
2,3-butanediol accumulation increases when potential is applied to the working electrode. HPLC measurement of 2,3 butanediol concentration in bioelectrochemical systems over time. Samples taken from working electrode chambers with electrodes poised at −0.3 V_SHE_ are shown in solid green and purple lines. Samples taken from bioelectrochemical systems disconnected from the potentiostat are shown in dashed green and purple lines. Samples from reactors containing cells with holo-PR are shown in green and those containing apo-PR are shown in purple. Each point represents the average of three replicates with standard error shown in error bars.

To determine whether acetoin reduction (as an electron sink) was necessary for inward electron transfer, we also performed experiments with strains lacking *bdh*. When Bdh was not expressed, no 2,3-butanediol was detectable in the bioreactors after 6 days, and cathodic current was significantly reduced, (p=0.02). The strain without Bdh generated ca. −11 µA when holo-PR was present (Figure 5), representing a ca. 32% reduction in electron transfer compared to the strain with Bdh. This indicates that full activity of the inward electron transfer system is dependent on acetoin reduction, although there is also some Bdh-independent electron transfer. This experiment also allows us to refine our comparisons between charge transfer and 2,3-butanediol accumulation. By subtracting the total charge transfer catalyzed by the strain without Bdh from the strain with Bdh, we can calculate the amount of charge transfer that is Bdh-dependent. We calculate that 0.03 mM of 2,3-butanediol accumulation is expected based on the amount of Bdh-dependent electron transfer. This amount of charge transfer agrees well with the 0.06 mM of electrode-dependent 2,3-butanediol accumulation that we observed.

**Figure 5.**
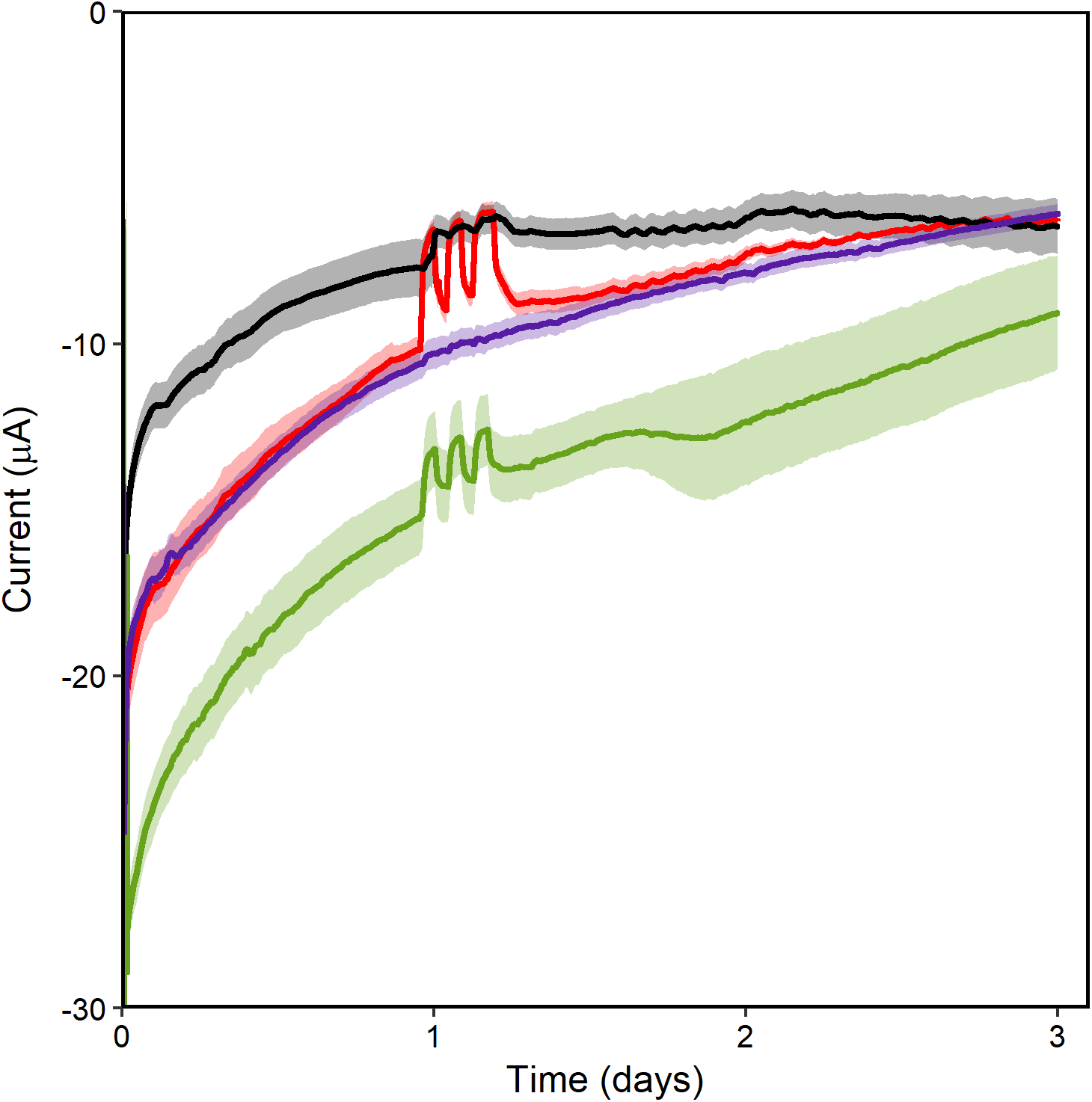
Both proteorhodopsin and butanediol dehydrogenase increase current. Current measured in bioelectrochemical systems containing cells expressing Bdh-PR with holo-PR are shown in green and those containing apo-PR are shown in purple. Current measured in reactors containing cells expressing only holo-PR are shown in red and those containing only apo-PR are shown in black. Current was measured in bioreactors with electrodes poised at −0.3 V_SHE_. Current shown begins at injection of acetoin (time 0). Each line represents the average of three replicates with standard error shown in transparent ribbons.

Electron uptake by the strain without Bdh was light-dependent when holo-PR was present, indicating that the Bdh-independent process requires PMF generation. This may indicate that the Bdh-independent process also relies on reverse activity of NADH dehydrogenases, although we do not yet know the eventual fate of electrons transferred to the cells without Bdh. No other typical metabolic byproducts of *S. oneidensis* MR-1 (e.g., acetate, formate) were detectable by our HPLC analysis.

### Hydrogen is an electron sink for inward electron transfer

A common question in previous work on microbial electrosynthesis is whether H_2_ mediates electron transfer, because it can be produced abiotically at the cathode (37). Because small amounts of hydrogen are difficult to measure and may be scavenged quickly by cells on the electrode, it has been challenging to rule out hydrogen as a mediator. We have attempted to do so using an approach similar to Deutzmann et al. (38), i.e., by using a hydrogenase mutant strain. The strain we used does not have the ability to use molecular hydrogen as an electron source or sink (35, 36, 39). To determine whether a hydrogen mediated pathway would function when hydrogenases were present, we also performed the experiments described above in a wild-type background. (Note: these experiments were performed without the initial oxic/anodic step, see Methods for details.) When hydrogenase activity was present, overall current increased while 2,3-butanediol accumulation decreased (Figure 6). This indicates that rather than acting as a mediator, H_2_ was an electron sink for *S. oneidensis* on the cathode. If hydrogen acted as a mediator in this system, we would expect a simultaneous increase in both current and 2,3-butanediol when hydrogenases were present. Our results indicate that cells with hydrogenases generated H_2_ and that when the pathway to hydrogen was cut off, electrons were directed more efficiently toward acetoin reduction.

**Figure 6.**
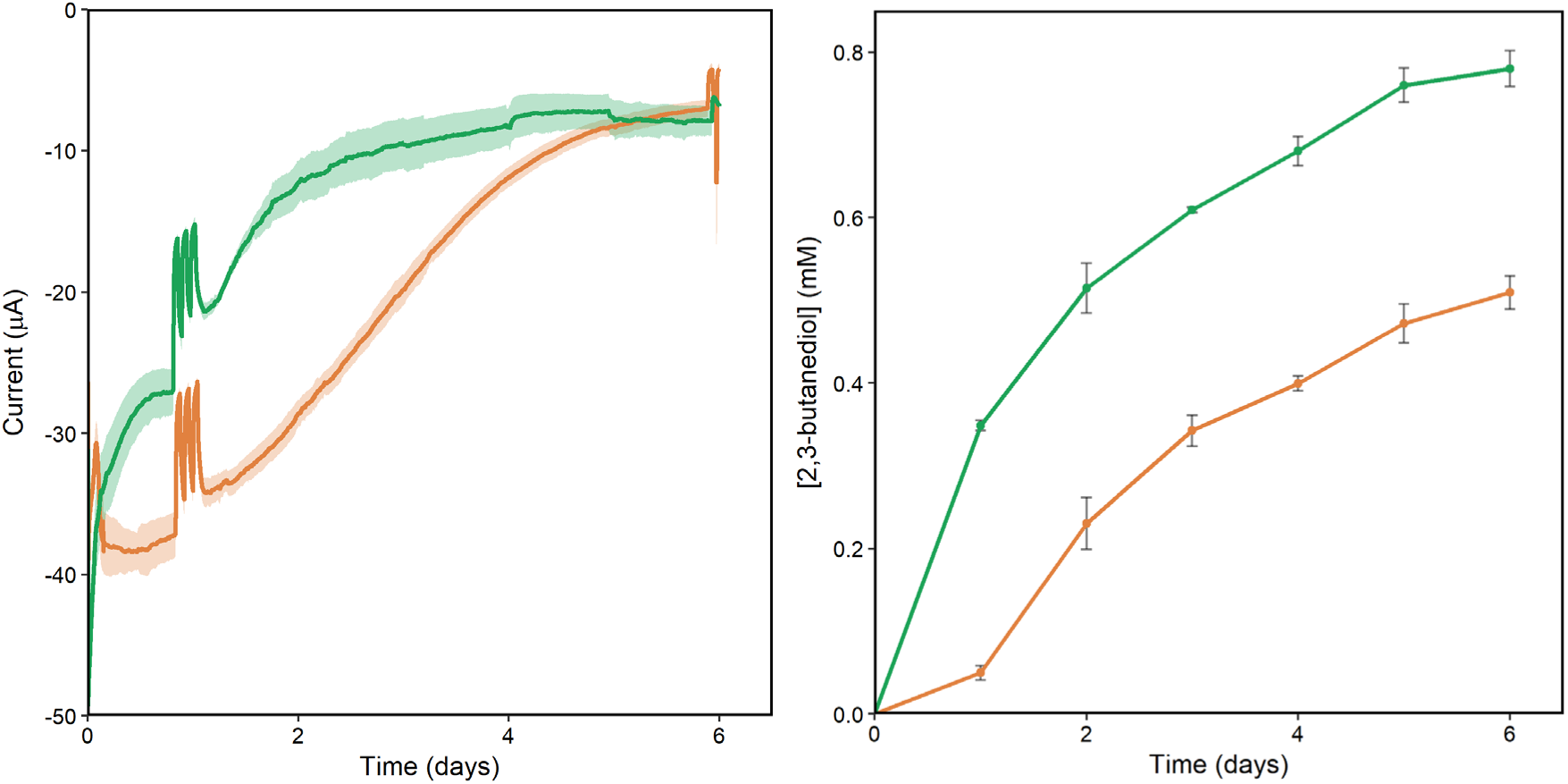
Removal of native hydrogenases decreases current and increases 2,3-butanediol production. (A) Current and (B) 2,3-butanediol accumulation observed in bioreactors containing cells expressing Bdh and holo-PR. The strain without hydrogenases is shown in dark green and the wild-type background is shown in orange. Data shown was measured beginning at injection of acetoin (time 0). Each point represents the average of three replicates with standard error shown in transparent ribbons or error bars.

### Comparison with previous work

Our results represent an advance in the field of microbial electrosynthesis because we have engineered a pathway to generate intracellular reducing power with a detailed understanding of the electron transfer mechanism. The transmembrane electron conduit connecting the quinol pool to electrodes has been very well characterized (21–23, 28, 40) and is known to be reversible (29,41). We hypothesized that we could take inward electron transfer in *S. oneidensis* MR-1 one step further by utilizing PMF to overcome the thermodynamic barrier of the quinol:NAD^+^ reduction and an excess of electron sink (Bdh/acetoin) to avoid over reduction of the NAD^+^:NADH pool. The results presented here support that hypothesis; we have shown that electron transfer to the target reaction is dependent on the electrode and on each of the modifications made to the strain. Further, the target reaction is NADH-dependent, indicating that we have successfully connected the electrode to the intracellular NADH pool.

While the system described here is one of the best understood inward electron transfer demonstrations to date, there is still more to learn. One critical knowledge gap is exactly how electrons are transferred from the quinol pool to NADH. The *S. oneidensis* MR-1 genome encodes four NADH:quinone oxidoreductases and any of them could be involved in inward electron transfer (42, 43). However, because we have observed dependence of inward electron transfer on PMF, we hypothesize that the proton-coupled NADH dehydrogenase, Nuo, is a major contributor to inward electron transfer. Future studies with Nuo deletion strains will enable us to elucidate the link between respiratory quinones and NADH in our system.

In this study, we have successfully driven a heterologous reduction reaction inside living bacteria using an extracellular electrode as the electron source. Rather than using native capabilities of acetogenic bacteria, we engineered a metal-reducing organism to reverse the flow of electrons in its respiratory chain. This represents a proof of concept for an engineered microbial electrosynthesis pathway, and with further development this platform can be used to upgrade bioproducts through electrofermentation or to fix CO_2_. Because electrical energy is one of the inputs to the system, the organic molecules produced also represent storage of electrical energy. This type of storage strategy will become essential as wind and solar power capacity increase. Wind and solar energy are intermittent sources; therefore, robust energy storage methods are critical. While significant improvements in engineered microbial electrosynthesis must still be made, a critical mass of researchers forming around this concept will help propel *Shewanella* based electrosynthesis toward real world application.

## Methods

### Bacterial strains, plasmids, and growth conditions

Strains and plasmids used in this study are listed in Table 1. *E. coli* strains were grown at 37°C and *S. oneidensis* strains at 30°C, both with shaking at 250 rpm. Strains were grown in LB medium (Miller, Accumedia) for assembly and initial verification of strains. Strains bearing pBBR1-MCS2-derived plasmids were grown with a final concentration of 50 µg/mL kanamycin. Pre-cultures for bioelectrochemical experiments were grown in M5 minimal medium: 1.29 mM K_2_HPO_4_, 1.65 mM KH_2_PO_4_, 7.87 mM NaCl, 1.70 mM NH_4_SO_4_, 475 μM MgSO_4_ ⋅ 7H_2_O, 10 mM HEPES, 0.01% (w/v) casamino acids, 1X Wolfe’s mineral solution (Al was not included), and 1X Wolfe’s vitamin solution (riboflavin was not included), pH adjusted to 7.2 with 5 M NaOH. M5 medium was supplemented with D,L-lactate to a final concentration of 20 mM. M5 medium with the following modifications was used in the working electrode chamber during bioelectrochemical experiments: 100 mM HEPES, no carbon source, no casamino acids, 1 µM riboflavin.

### Design and assembly of Bdh and Bdh-PR plasmids

The gene sequence encoding butanediol dehydrogenase in *Enterobacter cloacae* was downloaded from the NCBI gene database (NCBI Reference NC_014121.1). The codon usage was optimized for *S. oneidensis* MR-1 using JCAT Codon Adaptation Tool (www.jcat.de). A FLAG tag (44) was added immediately before the stop codon and the Salis lab RBS calculator (45) was used to design an optimized RBS for the Bdh-FLAG sequence. Sequences of 20 base pairs flanking the target SmaI restriction sites in the pBBR1-MCS2 sequence were added as flanking regions on the codon optimized RBS-Bdh-FLAG sequence for use in assembly. The entire sequence was submitted to Integrated DNA Technologies for synthesis as a gBlock gene fragment. The same general procedure was utilized to add the gene coding for proteorhodopsin to the plasmid containing *bdh*.

We isolated pBBR1-MCS2 plasmid DNA from *E. coli* using an E.Z.N.A plasmid DNA kit (Omega Bio-Tek). Prepared plasmid DNA was linearized using SmaI (New England Biolabs, Ipswich, MA) digestion for 4 hours at 25°C. The synthesized RBS-Bdh-FLAG insert was resuspended in water to a final concentration of 10 ng/µL and was assembled with linearized pBBR1-MCS2 plasmid using NEBuilder High Fidelity DNA assembly kit (New England Biolabs) using 50 ng of pBBR1-MCS2 and 100 ng of RBS-Bdh-FLAG insert. Assembled pBBR1-MCS2-Bdh was transformed into *E. coli* Mach1 chemically competent cells (Invitrogen). pBBR1-MCS2-Bdh-PR was prepared as above except pBBR1MCS2-Bdh was digested using NdeI and SpeI (New England Biolabs, Ipswich, MA) for 3 hours at 37°C prior to assembly with the synthesized PR gene. Transformants were initially screened via PCR using M13 forward and reverse primers and then sequenced (Sanger sequencing, RTSF Genomics Core, Michigan State University) to verify proper assembly and transformation. Verified Bdh and Bdh-PR plasmids were transformed into chemically competent *E. coli* WM3064 for use in conjugation with *S. oneidensis*. Conjugation was performed using a standard protocol for *S. oneidensis* MR-1 (46).

### Verifying expression by Western blot

Expression of Bdh and PR were verified via Western blot through use of FLAG tags added during gene synthesis. Cells were cultured in 5 mL of LB for 16 hours before 200 µL was centrifuged for 2 minutes at 10,000 rpm. Supernatant was removed, and cells were resuspended in 200 µL of a mixture of 1 mL Laemmli buffer, 20 µL of concentrated bromophenol blue (JT Baker, D29303) in 5X Laemmli buffer, and 10 µL of 1 M DTT. Cells were vortexed to mix and incubated at 95°C for 10 minutes. A mini-PROTEAN tetra cell electrophoresis chamber (Biorad, 1658005EDU) was loaded with 1X TGS buffer. Samples were vortexed a 5 µL was loaded onto a mini-protean TGX stain free gel (Biorad, 4568095) alongside 5 µL of Precision Plus ladder (Biorad, 1610376).

Samples were run at 100 V for 1.5 hours until dye front moved off the gel. The gel cassette was broken, and gel removed to 1X Transfer buffer (Biorad, 10026938). Proteins were then transferred to a nitrocellulose membrane (Biorad, 1704270) using a Biorad Turbo transfer system (Biorad, 1704150). The membrane was rinsed with 30 mL TBST buffer, this buffer was discarded, and the membrane was blocked using 50 mL of 3% BSA TBST buffer for 1 hour on an orbital shaker. Blocking solution was discarded and replaced with 30 mL of 3% BSA TBST buffer then 7.8 µL of 3.85 mg/mL anti-FLAG antibody (Sigma-Aldrich, F3165) was added. The membrane was then incubated for 16 hours at 4°C, on an orbital shaker.

The membrane was then rinsed for 5 minutes with TBST buffer three times. After rinsing 30 mL of 3% BSA TBST buffer with 0.0625 µL of anti-mouse antibody (Sigma-Aldrich, A9044) was added. The membrane was incubated for 1 hour at RT. Buffer was discarded and the membrane rinsed with TBST buffer for 5 minutes, three times. ECL Clarity chemiluminescence solution (Biorad, 1705061) was prepared by mixing 10 mL of peroxide solution with 10 mL of enhancer solution, then adding the entire volume to the membrane. The membrane was incubated for 5 minutes, the ECL solution was discarded and the membrane was imaged using a Kodak 4000R image station and Caresteam Molecular Imaging software.

### Bioelectrochemical system construction and operation

Bioelectrochemical measurements were performed in custom two-chambered bioreactors separated by a cation exchange membrane (Membranes International, CMI-7000S) cut in a 4.5 cm circle, to completely cover the 15 mm opening connecting working and counter chambers. Working electrodes were prepared from carbon felt (Alfa Aesar, 43200RF) cut into 50 × 25 mm rectangles and adhered to a titanium wire using carbon adhesive (Sigma-Aldrich, 09929-30G) and allowed to dry for 16 hours. Reference electrodes were prepared by oxidizing silver wires electrochemically in a dilute KCl solution and fixing them in saturated KCl-agar in a custom-made glass housing. The housing maintained ionic connection between the reference electrode and working chamber via a magnesia frit (Sigma-Aldrich, 31408-1EA). Counter electrodes were graphite rods 1/8” in diameter (Electron Microscopy Science, 07200) suspended in the counter chamber filled with PBS. Working chambers were filled with 140 mL of M5 medium (100 mM HEPES, no Casamino acids) prior to autoclaving. After autoclaving, 1.7 mL 100X vitamin stock, 1.7 mL Wolfe’s minerals (no Al), 0.17 mL 50 mg/mL kanamycin, and 0.85 mL 0.2 mM riboflavin were added to working chamber.

Reactors were connected to a potentiostat (VMP, BioLogic USA) and the working electrode poised at +0.4 V_SHE_. Current was measured every 1 second for the duration of the experiment. Current measurements were collected for at least 16 hours prior to inoculation. Reactors were inoculated with cultures grown in 50 mL M5 medium supplemented with 20 mM D,L-lactate. Cultures were grown in 250-mL flasks at 30°C for 17 hours shaking at 275 rpm. After 17 hours, 25 µL 20 mM all-*trans*-retinal (vitamin A aldehyde, Sigma-Aldrich, R2500), the essential proteorhodopsin cofactor (47), was added to designated flasks for functional PR testing to a final concentration of 10 µM and all flasks were returned to incubator, shaking, for 1 hour.

To achieve higher inoculum density without adjusting growth time and phase, two 50 mL culture volumes were prepared for each reactor. Absorbance at 600 nm was determined for each culture using a biophotomer (Eppendorf, D30) before the volume was transferred to a 50 mL conical tube (VWR, 89039-664) and centrifuged for 5 minutes at 10,000 rpm (Thermo Scientific ST8R; Rotor: 75005709). Supernatant was removed, and a second volume of cells was added to the conical tube containing the cell pellet before a second centrifugation step. Supernatant was removed, and the combined pellets were resuspended in 10 mL of M5 (100 mM HEPES).

Absorbance at 600 nm was determined for each prepared volume of cells before being standardized to OD_600_=3.6. The working chamber of each bioelectrochemical system was then inoculated with 9 mL of standardized cell suspension using an 18g needle (Beckton Dickson, 305196) and 10-mL syringe (Beckton Dickson, 302995). The working electrode was poised at +0.4 V_SHE_ for 6 hours, in the presence of ambient oxygen before the potential was changed to −0.3 V_SHE_, and N_2_ gas, 99.9% (Airgas) was bubbled into the reactors through a 0.2 µm filter. The rate of flow of nitrogen gas was observed using a bubbler attached to the gas outlet line from the reactors. The rate of gas flow was maintained so there was positive pressure against the water in the bubbler and bubble rate was kept constant between reactors. After 16 hours, sterile, anoxic acetoin solution was added to a final concentration of 10 mM.

Prior to inoculation, 0.93 m of green LED light strips (FAVOLCANO, 600 LEDs/5 m, 24 W/5 m, ≤6A /5 m) were attached to the exterior of the working chamber and switched on. Light cycling was performed at 24 hours post acetoin injection, lights were turned off for 1 hour followed by 1 hour on. Light cycles at 24 hours were repeated three times, after which lights were left on.

Samples were regularly removed from the bioreactors for determination of OD_600_ and HPLC analysis. A 2-mL sample was taken approximately every 24 hours using a 21g needle (Beckton Dickson, 305167) and 3-mL syringe (Beckton Dickson, 309657). One mL was used for determination of OD_600_. One mL was transferred to a micro-centrifuge tube (VWR, 20170-038) and frozen at −20°C until preparation for HPLC analysis. Experiments without a set potential on the working electrode were set up as above except, immediately after acetoin injection the potentiostat was disconnected from the working electrodes.

Experiments testing for the effect of hydrogenase activity (Figure 6) were performed as above expect as follows. Potential was set to −0.3 V_SHE_ during background data collection and no anodic potential was used. Oxygen removal using N_2_ gas was also started during background data collection and continued for the entire experiment. Cultures grown for reactor inoculation were standardized to the lowest observed OD_600_ after 18 hours of growth, instead of a target OD of 3.6. Finally, acetoin was injected two hours after inoculation once a stable base line current was achieved.

### HPLC Analysis

HPLC analysis was performed on a Shimadzu 20A HPLC, using an Aminex HPX-87H (BioRad, Hercules, CA) column with a Micro-guard Cation H^+^ guard column (BioRad, Hercules, CA) at 65°C. Compounds of interest were separated using a 0.6 mL/min flow rate, in 5 mM sulfuric acid with a 30-minute run time. Eluent was prepared by diluting a 50% HPLC-grade sulfuric acid solution (Fluka) in Milli-Q water and degassing the solution at 37°C for 3-5 days before use. Compounds of interest were detected by a refractive index detector (Shimadzu, RID-20A) maintained at 60°C. Samples were prepared by centrifuging 1-mL samples taken from the working electrode chambers for 10 minutes at 13,000 rpm in a microcentrifuge (Minispin Plus, Eppendorf) to remove cells. The supernatant was removed and transferred to a 2.0-mL glass HPLC vial (Vial: Restek, 21140; Cap: JG Finneran, 5395F09). Mixed standards of 2,3-butanediol and acetoin were prepared at concentrations of 1, 2, 5, 10, and 15 mM. Samples were maintained at 10°C by an auto-sampler (Shimadzu, SIL-20AHT) throughout analysis. Acetoin and 2,3-butanediol concentrations in the samples were determined using linear calibration curves based on the external standards.

## Data analysis

Analysis of HPLC and current data was performed using Rstudio using the following packages: ggplot2 (48), reshape2 (49), dplyr (50), and TTR (51).

## Acknowledgements

The authors thank Dr. Jeffrey Gralnick (University of Minnesota) for providing a plasmid containing the PR gene and Dr. N. Cecilia Martinez Gomez for helpful comments on the manuscript. This work was partially funded by NSF CAREER award 1750785. This work was also supported by the USDA National Institute of Food and Agriculture, Hatch project 1009805.

