## supplemental information for "Reversing an extracellular electron transfer pathway for electrode-driven NADH generation"

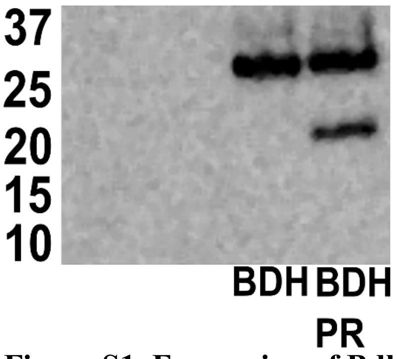


**Figure S1. Expression of Bdh and PR proteins.** FLAG tags were added to both genes during synthesis and used in detection of expression by Western blot using anti-flag antibodies.

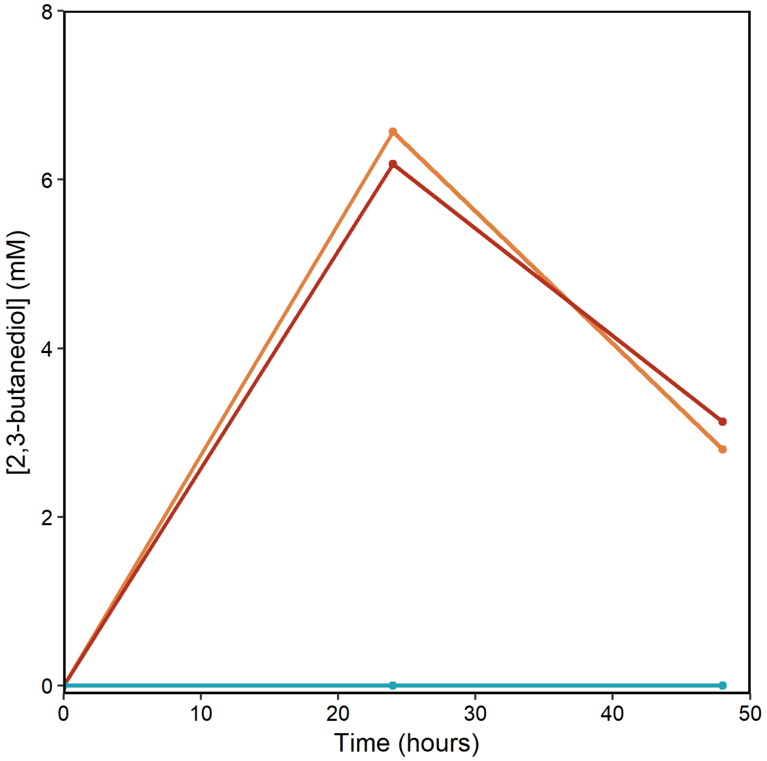


**Figure S2. 2,3-butanediol is produced aerobically by cells expressing heterologous Bdh.** Modified strains were grown in 50 mL LB with 15 mM acetoin in 250-mL flasks with 275 rpm shaking. Cells expressing Bdh alone are shown in red. Cells expressing both Bdh and PR are shown in orange, and cells expressing the vector alone are shown in teal. No detectable 2,3-butanediol was produced by cells lacking Bdh. Results are from single cultures of each strain.

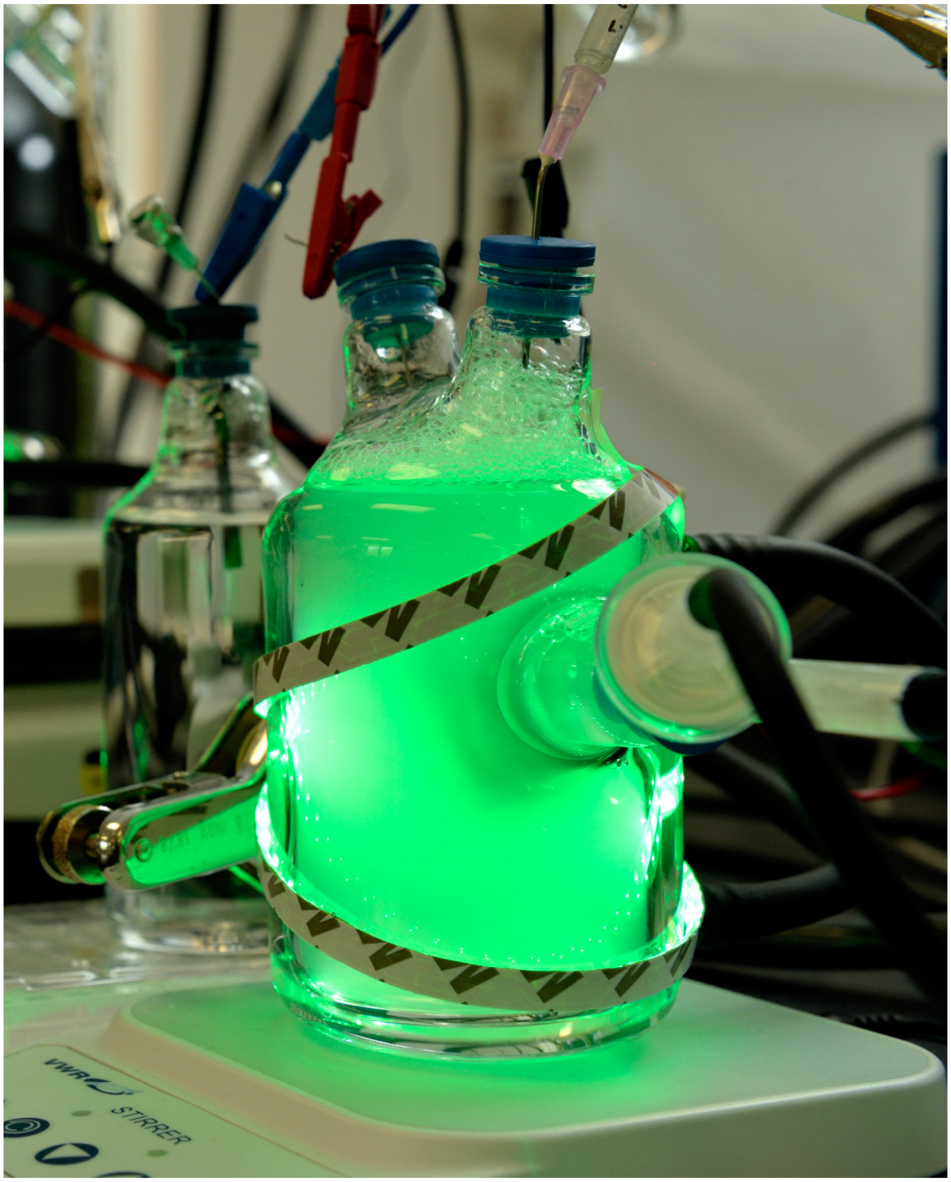


**Figure S3. Light is delivered to the system through the outer jacket of the working chamber.** Green LED light strips were fixed to the outside of the bioelectrochemical system. Above is a representative picture of this system.
